# G6b-B regulates an essential step in megakaryocyte maturation

**DOI:** 10.1101/2021.11.11.468226

**Authors:** Isabelle C. Becker, Zoltan Nagy, Georgi Manukjan, Melanie Haffner-Luntzer, Maximilian Englert, Tobias Heib, Timo Vögtle, Carina Gross, Richa Bharti, Sascha Dietrich, Kristina Mott, Johannes Heck, Anke Jeschke, Thorsten Schinke, Nicolas Schlegel, Tobias Heckel, David Stegner, Irina Pleines, Anita Ignatius, Harald Schulze, Bernhard Nieswandt

## Abstract

G6b-B is a megakaryocyte lineage-specific immunoreceptor tyrosine-based inhibition motif (ITIM)-containing receptor, essential for platelet homeostasis. Mice with a genomic deletion of the entire *Mpig6b* locus develop severe macrothrombocytopenia and myelofibrosis, which is reflected in humans with null-mutations in *MPIG6B*. The current model proposes that megakaryocytes lacking G6b-B develop normally, while proplatelet release is hampered, but the underlying molecular mechanism remains unclear. Here, we report on a spontaneous recessive single nucleotide mutation in C57BL/6 mice, localized within the intronic region of the *Mpig6b* locus that abolishes G6b-B expression and reproduces macrothrombocytopenia, myelofibrosis and osteosclerosis. As the mutation is based on a single nucleotide exchange, *Mpig6b^mut^* mice represent an ideal model to study the role of G6b-B. Megakaryocytes from these mice were smaller in size, displayed a less developed demarcation membrane system and reduced expression of receptors. RNA sequencing revealed a striking global reduction in the level of megakaryocyte-specific transcripts, in conjunction with decreased protein levels of the transcription factor GATA-1, and impaired thrombopoietin signaling. The reduced number of mature MKs in the bone marrow was corroborated on a newly developed *Mpig6b* null mouse strain. Our findings highlight an unexpected essential role of G6b-B in the early differentiation within the megakaryocytic lineage.

## Introduction

Megakaryocytes (MKs) are large, polyploid cells within the bone marrow (BM) that develop in a complex differentiation process. Mature MKs harbor granules and internal membrane systems, which are indispensable for the assembly and release of functional platelets.^1^ G6b-B has been identified as an essential regulator of platelet biogenesis.^2^ Deletion of the *Mpig6b* locus in mice results in macrothrombocytopenia and myelofibrosis,^2^ which was recently recapitulated in humans with disease-causing null-variants within *MPIG6B*.^3–7^ It was hypothesized that G6b-B affects a terminal step in platelet production,^2, 8^ however, mechanistic insights are yet lacking. By characterizing a spontaneous *Mpig6b* mutant together with a newly generated *Mpig6b*-null mouse line we provide unexpected experimental evidence that establishes G6b-B as a central regulator for MK maturation, which can explain the complex phenotype of these mice.

## Methods

Homozygous mutant mice carrying a spontaneously developed mutation in a splice acceptor site of *Mpig6b* (NM_001033221.3; c.404-1G>A) are referred to as *Mpig6b^mut^*. All methods are described in the supplementary methods. Whole exome sequencing data was submitted to NCBI SRA (BioProject-ID: PRJNA655378). RNA sequencing data are available at NCBI GEO (accession number: GSE155735).

## Results and Discussion

### Single nucleotide exchange in *Mpig6b* results in macrothrombocytopenia

In a breeding colony of C57BL/6 mice we identified individual animals with bleedings due to severe macrothrombocytopenia (**Figure 1a-b**). 10 generations of backcrossing led to isolation of a substrain presenting with a recessive trait. Using whole exome sequencing, we identified a homozygous single nucleotide exchange within *Mpig6b* (c.404-1G>A) in 10/10 mice (allele frequency: 1.0), resulting in abolished G6b-B protein expression in platelets (**Figure 1c-e, S1a-b**). In silico prediction unraveled the introduction of a splice acceptor site in intron 2-3 of *Mpig6b* leading to an out-of-frame shift transcript with multiple stop codons, expected to result in nonsense-mediated mRNA decay. Absence of G6b-B led to a reduction in surface expression levels of platelet membrane glycoproteins (GPs) (**Table S1**), infinite tail bleeding times, myelofibrosis, splenomegaly and additional osteosclerosis in female *Mpig6b^mut^* mice (**Figure S1c-f**).We thus identified a spontaneous single nucleotide mutation within *Mpig6b* resulting in a phenotype, which faithfully recapitulates *Mpig6b^−/−^* and *Mpig6b* diY/F mice.^2, 8, 9^

**Figure 1.**
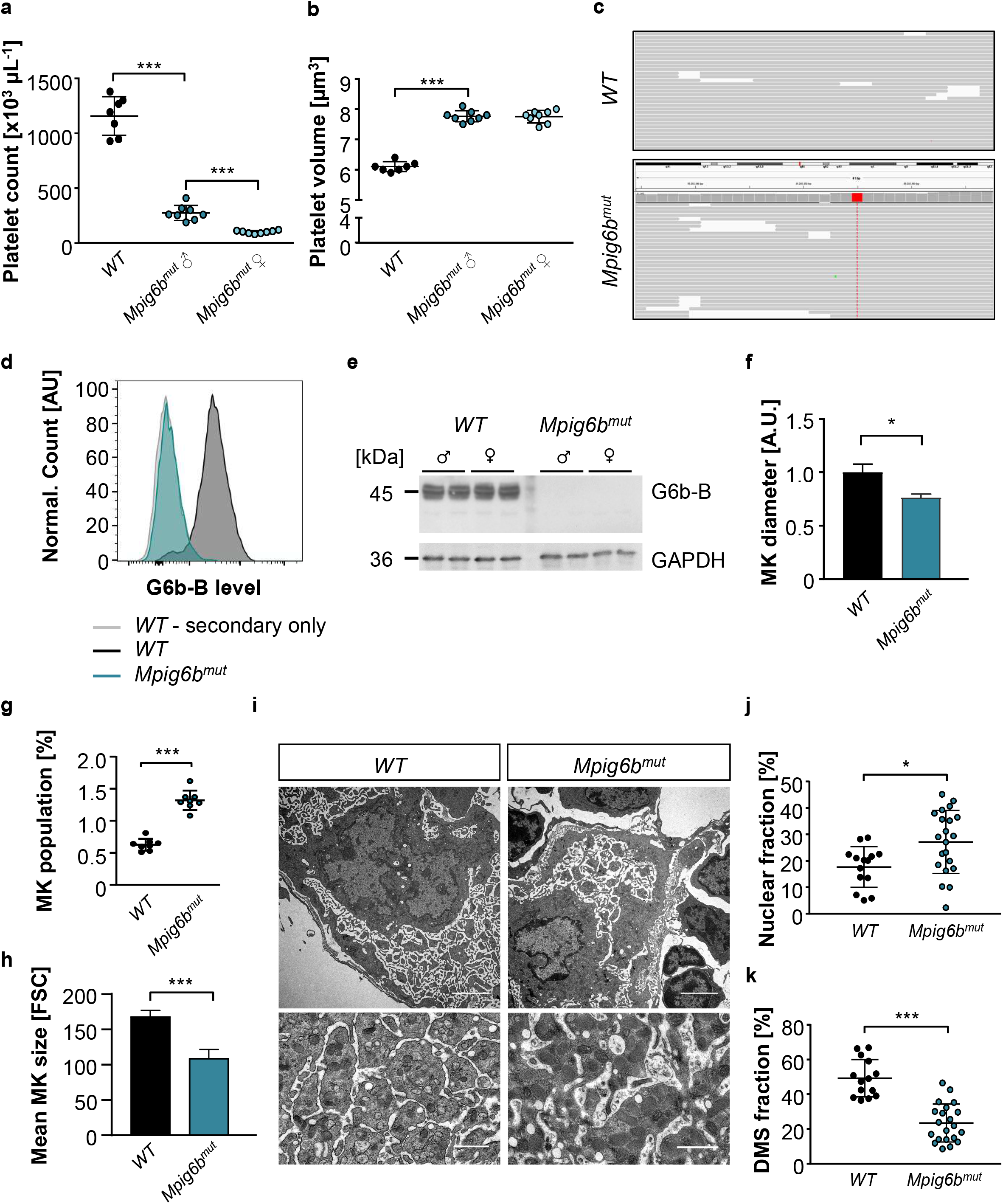
Single nucleotide mutation within *Mpig6b* results in severe macrothrombocytopenia and impaired MK maturation. Platelet count (**a**) and volume (**b**) in 10-week-old female and male *WT* and *Mpig6b^mut^* mice were assessed by an automated blood cell analyzer. Values are mean ± SD. Unpaired, two-tailed Student’s t-test. ***P < 0.001. (**c**) Identification of a mutation in a splice acceptor site of *Mpig6b* in *Mpig6b^mut^* mice by whole exome sequencing. The G>A single nucleotide exchange in *Mpig6b* was present in all reads in mutant, but not in *WT* mice. (**d-e**) Absence of G6b-B was validated in *Mpig6b^mut^* platelets by flow cytometry (**d**) and immunoblotting (C-terminal antibody) (**e**). (**f**) *WT* and *Mpig6b^mut^* MKs were matured in vitro in the presence of TPO and analyzed by brightfield microscopy. Mean MK diameter was determined manually using ImageJ software. At least 30 MKs per culture were analyzed. Values are mean ± SD (n = 3). Unpaired, two-tailed Student’s t-test. *P < 0.05. (**g**) The αIIbβ3-positive cell population in whole BM of *WT* or *Mpig6b^mut^* mice was analyzed by flow cytometry. Values are mean ± SD (n = 8). Unpaired, two-tailed Student’s t-test. ***P < 0.001. (**h**) Mean size of native MKs was analyzed ex vivo by flow cytometry. Values are mean ± SD (n = 4). Unpaired, two-tailed Student’s t-test. ***P < 0.001. (**i**) Demarcation membrane system (DMS) maturation in *WT* and *Mpig6b^mut^* BM MKs was visualized by TEM. Scale bars: 3 μm; insets: 1.5 μm. Nuclear (**j**) and DMS fraction (**k**) in relation to cell size were quantified manually using ImageJ software. At least 7 MKs per mouse were analyzed. Values are mean ± SD (n = 3). Unpaired, two-tailed Student’s t-test. *P < 0.05; ***P < 0.001.

Analyzing BM-derived MKs from *Mpig6b^mut^* mice in vitro, we found that the number of proplatelet-forming cells was significantly reduced compared to the wildtype (*WT*) **(Figure S2a-d)**. The cytoskeleton of proplatelet-forming *Mpig6b^mut^* MKs, however, appeared morphologically comparable. Intravital two-photon microscopy on the cranial BM revealed a high degree of MK fragmentation together with a low amount of circulating platelets in *Mpig6b^mut^* mice **(movie S1-3)**.

### Impaired maturation of *Mpig6b^mut^* MKs

The diameter of in vitro-matured *Mpig6b^mut^* MKs was significantly smaller (**Figure 1f**). When native BM cells were analyzed flow cytometrically, we observed an increased percentage of MKs in *Mpig6b^mut^* mice, also displaying a smaller size (**Figure 1g-h**). Transmission electron microscopy (TEM) analysis of BM MKs in situ revealed severely defective maturation of the demarcation membrane system (DMS) in *Mpig6b^mut^* mice (**Figure 2i-k**). We also observed increased neutrophil emperipolesis into mutant MKs in situ by TEM and in cryosections (**Figure S2e-f**). Ploidy analysis showed a significant rise in 2n megakaryoblasts, while the fractions of 16n and 32n MKs were marginally reduced (**Figure S2g-h**). Interestingly, the percentage of small- and medium-sized MKs was markedly increased, whereas the fraction of big and highly granular, side scatter^high^ (SSC^high^) MKs was severely reduced in the BM of *Mpig6b^mut^* mice (**Figure 2a-b**). The expression levels of prominent GPs positively correlated with MK size in *WT* animals, in which SSC^high^ MKs exhibited the highest GP surface abundance (**Figure S3a**). Notably, the overall GP surface expression in the entire MK population revealed a marked reduction for all GPs in *Mpig6b^mut^* mice (**Figure 2c**), due to the accumulation of immature MKs expressing lower levels of the respective GPs. These findings were recapitulated in in vitro-matured MKs (**Figure S3b-c**). The absence of a major block in MK ploidy levels together with reduced surface expression of GPIbα, α2 integrin and GPVI are coherent with data from *Mpig6b^−/−^* and *Mpig6b^fl/fl^;Pf4-Cre^+^* mice.^2, 10^ Although these findings exclude a direct role for G6b-B in endomitosis, they point to a central function in regulating MK maturation. Interestingly, this resembles other mouse lines, where reduced MK maturation is associated with normal ploidy, including *Gfi1b^−/−^* 11, *Nfe2^−/−^* mice^12^, or *Rhoa^fl/fl^;Cdc42^fl/fl^;Pf4-Cre^+^* mice.^13^ The accumulation of immature MKs can lead to myelofibrosis, osteosclerosis, or both, as reported for *Gata-1^low^* or *Nfe2^−/−^* mouse lines,^14–16^ and might explain the phenotypes in G6b-B null mice.

**Figure 2.**
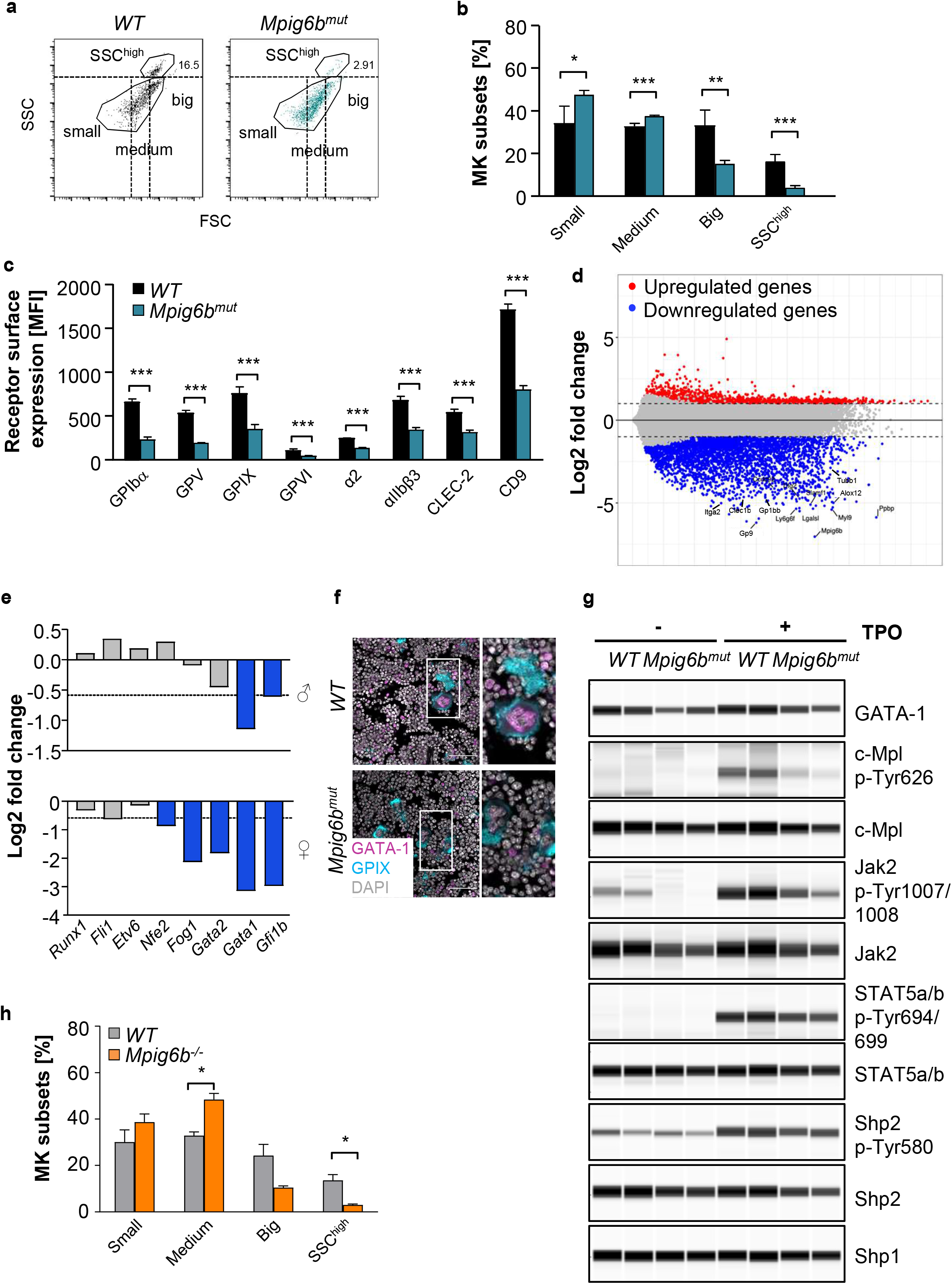
Maturation block in *Mpig6b^mut^* MKs involves reduced gene expression and TPO signaling. (**a, b**) Mean size distribution of *WT* and *Mpig6b^mut^* MKs was analyzed by flow cytometry. (**a**) Dot plots depicting the proportion of SSC^high^ MKs and the delineation between small, medium and big MKs. (**b**) Values are mean ± SD (n = 6). Unpaired, two-tailed Student’s t-test. *P < 0.05; **P < 0.01; ***P < 0.001. (**c**) Mean surface receptor expression on the whole MK population derived from *WT* and *Mpig6b^mut^* mice. Values are mean ± SD (n = 4). Unpaired, two-tailed Student’s t-test. ***P < 0.001. (**d**) Volcano plot showing up- and downregulation of genes in native *Mpig6b^mut^* MKs derived from female mice compared to female *WT* controls. Black lines point towards downregulated MK-specific genes (e.g. *Tubb1, Clec1b, Gp1bb*). (**e**) Up- and downregulation of MK differentiation-related transcription factors in native MKs from male and female *Mpig6b^mut^* mice compared to the respective control (n = 4). Only values with a log2 fold change <0.5849 (dotted line) were considered downregulated. (**f**) Immunostainings of femora cryosections visualizing GATA-1 expression in *WT* and *Mpig6b^mut^* MKs in situ. Scale bars: 50 μm. *Runx1*: Runt-related transcriptions factor 1; *Fli1*: Friend leukemia integration 1 transcription factor; *Etv6*: ETS variant transcription factor 6; *Fog1*: Friend of GATA-1; *Gfi1b*: growth factor-independent 1B transcriptional repressor. (**g**) Phosphorylation and/or total expression levels of GATA-1, c-Mpl, Jak2, STAT5a/b, Shp1 and Shp2 in TPO-stimulated *WT* and *Mpig6b^mut^* MKs were visualized using an automated quantitative capillary-based immunoassay platform; Jess (ProteinSimple). (**h**) Mean size distribution of *WT* and *Mpig6b^−/−^* MKs was analyzed by flow cytometry. Values are mean ± SD (n = 2). Unpaired, two-tailed Student’s t-test. *P < 0.05.

### Defective MK-specific gene expression in *Mpig6b^mut^* mice

Next, we performed bulk RNA sequencing on purified, native BM MKs from young adult mice, not yet displaying myelofibrosis. The transcriptome signature of *Mpig6b^mut^* MKs revealed a striking decrease in a plethora of MK-specific transcripts, including *Tubb1*, *Myl9* or *Gp1ba* (**Figure 2d**). mRNA levels of transcription factors GATA-1and Gfi1b, which are indispensable for MK differentiation,^17, 18^ were significantly less abundant in *Mpig6b^mut^* MKs (**Figure 2e**). qPCR analysis revealed that transcript levels of *Itgb3*, *Tubb1* and *Gata1* were also significantly reduced when mutant MKs were in vitro-differentiated **(Figure S3d)**. In addition, GATA1 protein levels were reduced in both native MKs in situ **(Figure 2f)**, as well as in in vitro-differentiated *Mpig6b^mut^* MKs **(Figure 2g, S3e)**. Our RNA-sequencing data strongly implies that loss of G6b-B hampers early transcriptional pathways within the megakaryocytic lineage, as reflected by downregulation of many MK-specific genes in primary MKs.

### Defective TPO-signaling in *Mpig6b^mut^* MKs

*Mpig6b^mut^* mice exhibited elevated thrombopoietin (TPO) plasma levels (**Figure S3e**), as expected in response to low platelet counts. We thus investigated the total MK protein and phosphorylation levels of proteins, which are crucial to TPO signaling,^10, 19^ Protein levels of c-Mpl, Jak2 and Shp2 were decreased in *Mpig6b^mut^* MKs compared to *WT* MKs, while STAT5a/b were not significantly altered. Moreover, c-Mpl, Jak2, STAT5a/b and Shp2 phosphorylation was markedly reduced (**Figure 2g, S3f**). Our findings thus provide evidence that loss of G6b-B disturbs TPO-based signaling events in MKs and point to a critical role of G6b-B in regulating relevant signaling pathways driving early MK maturation.

### *Mpig6b^−/−^* mice display less mature MKs

To corroborate that our findings reflect the role of G6b-B and are not the consequence of an unlikely co-segregating mutation, we generated a novel *Mpig6b^−/−^* mouse model (**Figure S4a-d**). MKs from this novel *Mpig6b^−/−^* mice displayed an increase in the percentage of small- and medium-sized MKs and a marked decrease in big and SSC^high^ MKs (**Figure 2h**), thus confirming that lack of G6b-B results in a MK maturation defect.

Patients with disease-causing variants within *MPIG6B* present(ed) with congenital macrothrombocytopenia, mild-to-moderate bleeding diathesis, focal myelofibrosis and atypical MKs, however the underlying causes of the disease have remained unclear.^3–7^ Our results demonstrate a previously unrecognized key function of G6b-B in MK maturation by regulating cell size, DMS development, receptor levels and gene expression. These findings thus challenge the previous concept, which proposed a rather unaffected development of G6b-B deficient MKs^2, 4, 8, 20^, and provide an unexpected mechanistic explanation for the severe macrothrombocytopenia in *Mpig6b^mut^* mice, as a direct consequence of an overall MK maturation defect. We observed an increased number of immature MKs in the bone marrow, which might help to better understand the cause of myelofibrosis in G6b-B null mice and patients, as an accumulation of immature MKs has been described as the main driver of this complex disease.^21–25^

## Acknowledgements

We thank Stefanie Hartmann, Sylvia Hengst, Daniela Naumann and Mariola Dragan for excellent technical assistance and the microscopy platform of the Bioimaging Center Würzburg for providing technical infrastructure and support. We thank Prof. Dr. Yotis Senis (Strasbourg, France) for providing anti-G6b-B antibodies. We also thank the Core Unit SysMed at the University of Würzburg for excellent technical support and RNA-seq data generation.

## Funding

Deutsche Forschungsgemeinschaft (DFG, German Research Foundation) (Project number 374031971 - TRR 240/project A01 to BN and project A03 to HS, and grant NI 556/11-2 to BN)

IZKF at the University of Würzburg (project Z-6)

European Union (EFRE - Europäischer Fonds für regionale Entwicklung, Bavaria)

German Excellence Initiative to the Graduate School of Life Sciences, University of Würzburg (to IB and ZN)

## Author contributions

Conceptualization: ICB, ZN,GM, HS, BN; Methodology: MHL, RB, SD; Software: TH, RB, SD; Investigation: ICB, ZN, GM, MHL, ME, TH, TV, CG, JH, AJ; Resources: TS, NS, TH, DS, HS, BN; Supervision: IP, AI, HS, BN; Writing - original draft: ICB, ZN, GM, HS, BN; Writing— review & editing: ICB, ZN, GM, MHL, IP, HS, BN

## Declaration of Interests

The authors declare no competing interests.

## Data availability

Whole exome sequencing data is available under the BioProject-ID PRJNA655378 (https://www.ncbi.nlm.nih.gov/bioproject/). Sequencing data are available at NCBI GEO (http://www.ncbi.nlm.nih.gov/geo) under the accession number GSE155735.

## Notes

### Competing Interest Statement

The authors have declared no competing interest.

