## Supplemental Material for "G6b-B regulates an essential step in megakaryocyte maturation"

### Supplementary Methods

#### Mice

In this study, 3- to 12-week-old mice were used for all experiments if not stated otherwise. All animal studies were approved by the district government of Lower Franconia (Bezirksregierung Unterfranken; AZ: 2-351; 2-4). A novel *Mpig6b*$^{-/-}$ mouse strain lacking exon1 and part of exon 2 resulting in a less than 500 base pair chromosomal deletion has been generated by CRISPR/Cas9-based Extreme Genome Editing technology (Biocytogen Inc., China).

#### Whole exome sequencing

DNA from eight mutant and eight wildtype mice was purified using the GeneJET Genomic DNA Purification Kit (Thermo Scientific) after retro-orbital bleeding. DNA quality was assured by high-resolution electrophoresis on a Bioanalyzer (Agilent). Whole Exome Sequencing of 50 ng mouse genomic DNA was performed according to Agilent's protocol for the SureSelect QXT reagent kit and SureSelect XT Mouse All Exon plus capture probes for Illumina sequencing (49.6 Mb capture spanning 221,784 exons and 24,306 genes). 2 × 75 bp paired-end sequencing reads were generated on the NextSeq-500 platform. Reads were processed using FastQC[1] (v0.11.51) for assessing read quality, amount of duplicates and presence of adapter sequences. Further, the processed sequences were aligned to Mouse Build 38 patch release 5 from GENCODE[2] (Burrows–Wheeler Aligner) using default parameters. Quality score recalibration was performed to prevent false-positive SNV calls and obtain accurate scores on SNV calls at the end of sequencing reads. Finally, SnpEff (v4.3p) was used to annotate the variants and was evaluated using GQ, DP and AD parameters indicating the confidence (GQ) of total informative reads at the site (DP) and number of alternative allele at the site (AD). The intronic missense mutation c.404-1G>A within *Mpig6b* was validated by sanger sequencing (n = 2). Sequencing data was submitted to NCBI SRA and is available under the BioProject-ID PRJNA655378.

#### Blood parameters

Platelet count, size and basic blood parameters were obtained in EDTA-anticoagulated blood using an automated cell counter (ScilVet, scil animal care company GmbH).[3]

#### Platelet preparation

Mice were anaesthetized using isoflurane and retro-orbitally bled into heparin (20 U/mL, Ratiopharm) or sodium citrate. After centrifugation at 800 rpm (twice), platelet-rich plasma was obtained, supplemented with 2 µL/mL apyrase (0.02 U/mL; A6410, Sigma-Aldrich) and 5 µL/mLPGI$_2$ (0.1 µg/mL; P6188, Sigma-Aldrich) and centrifuged again at 2800 rpm for 5 min.

Platelets were washed twice using Tyrode's-HEPES buffer w/o $Ca^{2+}$ (134 mM NaCl, 0.34 mM $Na_2HPO_4$, 2.9 mM KCl, 12 mM $NaHCO_3$, 5 mM HEPES, 5 mM glucose, 0.35% BSA, pH 7.4) and afterwards allowed to rest for 30 min at 37°C in Tyrode's buffer containing 0.02 U/mL apyrase.

**Transmission electron microscopy (TEM)**

For analysis of BM MKs, femora were isolated, segmented into 3 pieces using a scalpel and fixed using Karnovsky buffer (2% PFA, 2.5% glutaraldehyde in 0.1 M cacodylate buffer) overnight at 4°C. Afterwards, bones were incubated in 10% EDTA in PBS for 5 days for decalcification, followed by fixation of fatty components using 2% osmium tetroxide in 50 mM cacodylate buffer (pH 7.2). After fixation, samples were stained with 0.5% aqueous uranyl acetate, dehydrated via incubation in a graded ethanol series and embedded in Epon 812. Upon staining with 2% uranyl acetate (in 100% ethanol) and lead citrate, ultra-thin sections were imaged at a JEM-2100 (JEOL). Nuclear and DMS fraction were calculated by manually measuring DMS and nucleus size in relation to cell size using ImageJ Software.

**Flow cytometry on platelets**

Expression levels of different GPs were assessed in PBS-diluted blood using the respective fluorophore-conjugated antibodies by flow cytometry at a FACSCalibur or Celesta (BD Biosciences). G6b-B expression levels were analyzed on PFA-fixed, washed platelets using a FITC-conjugated anti-rat secondary antibody[4] (kindly provided by Prof. Dr. Yotis Senis, University of Strasbourg, France).

**Tail bleeding**

2 mm of the tail tip of anaesthetized mice was cut off with a scalpel. Bleeding time was assessed by gently absorbing blood on filter paper every 20 s until cessation of bleeding. After 20 min, experiments were stopped and mice were sacrificed. Fisher's exact test was used to analyze differences between definite and infinite bleeding.

**Histology**

Paraformaldehyde (PFA-)fixed, dehydrated and paraffin-embedded femora and spleens were cut into 3 µm sections, dried at 37°C overnight and stained with hematoxylin (MHS32, Sigma-Aldrich) and eosin (318906, Sigma-Aldrich). For visualization of reticulin fibers, sections were acquired as mentioned above and stained using the Reticulin Stain Kit (Polysciences) according to the manufacturer's suggestions. Image acquisition was performed at an inverted light microscope (Leica DMI4000B).

**Enzyme-linked immunosorbent assay (ELISA)**

Mice were bled up to 700 µl into 70 µl sodium citrate and blood was centrifuged at 2800 rpm, followed by retrieval of plasma by centrifugation at 14000 rpm. Plasma was immediately stored at -80°C until further processing. TPO levels were determined using a mouse TPO quantikine ELISA set (MTP00, R&D Systems) according to the manufacturer's protocol. Samples were analyzed at a Multiskan Ascent (96/384) plate reader (MTX Lab Systems) at 450 nm with wavelength correction (570 nm).

**Isolation of native MKs**

To obtain native MKs, BM was isolated from tibiae and femora by centrifugation and MKs and precursors were isolated using an anti-CD61 antibody coupled to magnetic beads (Miltenyi Biotec). Immediately afterwards, cells were subjected to a BSA density gradient separation and centrifuged at 300 g for 5 min.

**RNA sequencing**

RNA quality was checked using a 2100 Bioanalyzer with the RNA 6000 Pico kit (Agilent Technologies). The RIN for all samples was ~8. DNA libraries suitable for sequencing were prepared from 50 ng of total RNA with oligo-dT capture beads for poly-A-mRNA enrichment using the TruSeq Stranded mRNA Library Preparation Kit (Illumina). Sequencing was performed on the NextSeq-500 platform (Illumina) in single-end mode with 75 nt read length. Sequencing reads were trimmed for adapter sequence, mapped to the mouse genome (GRCm38.p6) with STAR and gene level based read counts were generated with featureCounts using the RefSeq annotation. The count output was utilized to identify differentially expressed genes using DESeq2[5] with log fold change shrinkage using betaPrior[6]. Clusterprofiler was used to perform gene set enrichment analysis for KEGG and GO pathways and to make an enrichment map for all of the pathways with a q-value < 0.01.[7] Deconvolution of relative immune cell composition on mouse expression profiles was performed with the ImmuCC algorithm.[8] Sequencing data are available at NCBI GEO (http://www.ncbi.nlm.nih.gov/geo) under the accession number GSE155735.

**MK preparation**

Mice were sacrificed and tibiae and femora were isolated. BM was obtained by centrifugation 2500g for 40s as recently described.[9] Megakaryocytic precursors were separated by lineage depletion using an antibody mixture (Lineage depletion panel, 133307, Biolegend) and magnetic beads (CD4 untouched, Miltenyi Biotec). Cells were incubated in TPO (50 ng/mL) alone (for qPCR and immunoblotting) or TPO- and hirudin- (100 U/mL) (rHirudin, Hyphen Biomed, RE120A) containing medium (Dulbecco's Modified Eagle Medium; Thermo Fisher

Scientific). After 72h MKs were enriched by a BSA density gradient and either used directly or further incubated in the presence of TPO and rHirudin for 24h to assess proplatelet formation. Cells were imaged at an automated microscope (EVOS M5000, ThermoFisher Scientific) and mean cell size was measured using ImageJ Software (NIH).

**Cryosectioning and immunofluorescence staining**

Mice were sacrificed and femora were isolated and fixed in 4% PFA containing 5 mM sucrose for 1h at RT. Organs were transferred into 10% sucrose in PBS and a sucrose gradient was performed over 3 days. Bones were stored in 30% sucrose until further processing. Femora were cut at 10 µm and transferred onto slides using a tape-transfer system (Kawamoto) at a Cryostat (Leica) and rehydrated in PHEM for 15 min. Sections were fixed in 4% PFA in PHEM, blocked using 3% goat serum and stained using antibodies against CD105 (120402 (MJ7/18), Biolegend), Ly6G (clone 1A8), and GATA-1 (#3535, Cell Signaling). MKs were visualized using directly labeled antibody derivatives against GPIX. After washing, slides were mounted using 4′,6-diamidino-2-phenylindole (DAPI)-containing Fluoroshield (Sigma-Aldrich). Image acquisition was performed at a confocal microscope using a 25x objective (Leica TCS SP8). MK numbers and emperipolesis were quantified manually using ImageJ Software (NIH).

**Immunofluorescence staining of proplatelet-forming MKs**

Proplatelet-forming MKs (approximately 300 cells per mouse) were analyzed the following day at a Zeiss Primovert. For immunofluorescence staining, glass slides were coated using 0.01% Poly-L-Lysine (P8920, Sigma-Aldrich), MKs were diluted in platelet buffer (10 mM HEPES, 140 mM NaCl, 3 mM KCl, 0.5 mM $MgCl_2$, 5 mM $NaHCO_3$, 10 mM glucose) and spun onto the slides at 800 rpm for 4 min. Cells were fixed, antibody-binding was blocked using 3% BSA and the cells were permeabilized with 0.1% TritonX100. MK cytoskeleton was stained using an anti-α-tubulin AF488-conjugated antibody (3.33 mg $mL^{-1}$, 322588 (B-5-1-2), Invitrogen), phalloidin-Atto647N (170 nM, 65906, Fluka). Cells were mounted using DAPI-containing Fluoroshield. Images were acquired at a Leica TCS SP8 microscope and analyzed using ImageJ software (NIH).

**Flow cytometry on MKs**

BM was centrifuged out of the bones as described above and flushed through a 70 µm cell strainer. For analysis of GP expression on BM MKs, unspecific binding was blocked on in vitro-matured MKs or in whole BM using 0.02 mg/mL anti-FcγR antibody (2.4G2, BD Pharmingen). Cells were washed, resuspended in CATCH buffer (25 mM HEPES, 3 mM EDTA, 3.5% BSA in PBS) and stained for 20 min on ice using FITC-conjugated antibodies against GPIb, GPIX, GPV, GPVI, CLEC2, αIIbβ3, α2 and CD9. MKs were distinguished from other BM cells by concomitant staining using an anti-αIIbβ3-antibody (10 µg/mL, clone MWReg30). Cells were analyzed on a FACSCelesta.

**MK ploidy**

MK ploidy was determined in whole BM after centrifugation. MKs were stained using a fluorescein isothiocyanate-conjugated anti-αIIbβ3 antibody (10 mg/mL, clone MWReg30) after blockade of unspecific binding by incubation with 0.02 mg/mL anti-FcgR antibody (2.4G2, BD Pharmingen). Subsequently, cells were fixed and permeabilized followed by labeling of DNA using 50 µg/mL propidium iodide (P3561, Invitrogen) containing 100 µg/mL RNaseA (EN0202, Fermentas) in PBS. Ploidy was assessed by flow cytometry on a FACSCalibur (BD Biosciences) and analysis was done using FlowJo software (Version 10; Tree Star Inc.).

**qPCR**

TPO-conditioned MKs or native MKs were washed in PBS once and lysed using Trizol reagent (Invitrogen). RNA was extracted using the RNeasy Mini Extraction Kit (Qiagen) according to the manufacturers' protocol. Using a staring amount of 500 µg RNA, cDNA was generated by reverse transcription with the iScript Select cDNA Synthesis Kit (Bio-Rad) following the manufacturers' recommendations with random primers. The iTaq Universal SYBR Green Supermix (Bio-Rad) was utilized for quantitative real-time PCR. *Sdha* and *Actb* served as housekeeping genes for the calculation of relative expression by the ΔΔCt method.

**Immunoblotting**

Platelets ($1 \times 10^6$ $mL^{-1}$) were prepared as described above and lysed for 30 min on ice. Supernatant was stored at -20°C until further analysis. Proteins were separated by SDS-PAGE and onto PVDF membranes. Membranes were blocked using Blueblock PF (Serva Electrophoresis GmbH) and probed with an anti-G6b-B antibody[4]. GAPDH (1 µg $mL^{-1}$, G5262, Sigma-Aldrich) served as a loading control. Horseradish-peroxidase conjugated secondary antibodies (0.33 µg/mL) and enhanced chemiluminescence solution (JM-K820-500, MoBiTec) were used to detect proteins.

For analysis of TPO signaling, in vitro-differentiated MKs were seeded into 24-well-plates at a density of $3.5 \times 10^5$ cells, starved for 4h in DMEM containing 0.5% FCS followed by stimulation with 50 ng $mL^{-1}$ TPO for 10 min. Cells were centrifuged at 2800 rpm for 1 min and immediately lysed on ice using 1x RIPA containing 1x Halt protease and phosphatase inhibitor (78440, ThermoFisher Scientific). Phospho- and/or total levels of Jak2, STAT3, STAT5a/b, Shp1, Shp2, GATA-1 and c-Mpl were assessed with an automated capillary-based immunoassay platform (Jess, ProteinSimple) according to the manufacturers' recommendations. *WT* and *Mpig6b*$^{mut}$ MK lysates were analyzed at equal protein concentrations. Separation matrix loading time was set to 200 seconds, stacking matrix loading time to 15 seconds, sample loading time to 9 seconds, separation time to 31 minutes, separation voltage to 375 volts, antibody diluent time to 5 minutes, primary antibody incubation time to 90 minutes and secondary antibody incubation time to 30 minutes. For recording of the chemiluminescent signal a High Dynamic Range (HDR) profile was used. Stat5a/b was considered as loading control since the expression of these transcription factors was not significantly altered in our RNA-Seq dataset.

**Data analysis**

The presented results are mean ± standard deviation (SD). Data distribution was analyzed using the Shapiro-Wilk-test and differences between control and mutant mice were statistically analyzed using unpaired, two-tailed Student's t-test, one- or two-way ANOVA. Tukey or Sidak's post-hoc test was used for multiple comparisons. P-values < 0.05 were considered as statistically significant *P < 0.05; **P < 0.01; ***P < 0.001.

**Figure S1. Increased bleeding time, myelofibrosis, splenomegaly and osteosclerosis in *Mpig6b$^{mut}$* mice.** (**a**) Identification of a mutation in a splice acceptor site of *Mpig6b* in *Mpig6b$^{mut}$* mice by whole exome sequencing. Two other SNPs were identified in *Git1* and *Prmd1,* but were classified as non-pathogenic with allele frequencies of 0.5 and 0.8, respectively, being insufficient to explain the homozygous trait with full penetrance. (**b**) Mutation (c.404-1G>A) was confirmed by Sanger sequencing. (**c**) Hemostasis in female and male *Mpig6b$^{mut}$* mice and the respective *WT* controls was analyzed in a tail bleeding assay on filter paper. Differences between definite and infinite bleeding were analyzed using a Fisher's exact test (n = 10). (**d**) Reticulin fibers within BM paraffin sections of 6-week-old *WT* and *Mpig6b$^{mut}$* mice were silver-stained. Nuclei were counterstained using nuclear fast red. Scale bars: 75 µm. (**e**) Representative images of spleens from male and female *WT* and *Mpig6b$^{mut}$* mice. (**f**) HE-stained BM paraffin sections of 10-week-old female and male WT and *Mpig6b$^{mut}$* mice reveal sex-specific osteosclerosis. Scale bars: 100 µm.

**Figure S2. Reduced proplatelet formation and increased neutrophil emperipolesis in *Mpig6b$^{mut}$* mice.** (**a**) Proplatelet-forming and round MKs were imaged using brightfield microscopy. Scale bars: 100 µm (**b**) Percentage of proplatelet-forming MKs was counted manually at a brightfield microscope. An average of five visual fields per well is shown. Values are mean ± SD (n = 4). Unpaired, two-tailed student's t-test. **P < 0.01. (**c**) F-actin and α-tubulin distribution in proplatelet-forming *WT* and *Mpig6b$^{mut}$* MKs was visualized by confocal microscopy (Leica TCS SP8, 40x objective). Nuclei were visualized using DAPI. Scale bars: 40 µm; insets: 10 µm. (**d**) Proplatelet tip size and numbers were determined manually using ImageJ software. Values are mean ± SD (n = 3). Unpaired, two-tailed student's t-test. ***P < 0.001. (**e**) Neutrophil emperipolesis into *Mpig6b$^{mut}$* MKs was visualized by TEM (scale bar: 3 µm) and in whole femora cryosections derived from male *WT* and *Mpig6b$^{mut}$* mice stained for GPIX and Ly6G. (**f**) Neutrophil emperipolesis was quantified from cryosections. Values are mean ± SD (n = 3). Unpaired, two-tailed Student's t-test. **P < 0.01. (**g, h**) Histogram and ploidy distribution of native BM MKs derived from 6-week-old *WT* and *Mpig6b$^{mut}$* mice. Values are mean ± SD (n = 6). Unpaired, two-tailed Student's t-test. *P < 0.05; ***P < 0.001.

**Figure S3. Defective maturation of *Mpig6b$^{mut}$* MKs in vitro.** (**a**) Upregulation of MK-specific surface receptors on *WT* BM MKs. Cells were distinguished according to their size as determined in Figure 2a. Values are mean ± SD (n = 4). Mean surface receptor expression (**b**) and size distribution (**c**) on in vitro-differentiated MKs derived from *WT* and *Mpig6b$^{mut}$* mice was analyzed by flow cytometry. Values are mean ± SD (n = 4). Unpaired, two-tailed Student's t-test. *P < 0.05; **P < 0.01; ***P < 0.001. (**d**) Relative mRNA expression of *Itgb3*, *Tubb1* and *Gata1* was analyzed by qPCR using in vitro-differentiated MKs. Values are mean ± SD (n = 3).

Unpaired, two-tailed Student's t-test. *P < 0.05; ***P < 0.001. (**e**) Plasma TPO levels were determined using an enzyme-linked immunosorbent assay. Values are mean ± SD (n = 5). Unpaired, two-tailed Student's t-test. ***P < 0.001. (**f**) Quantification of phosphorylation and/or total levels of GATA-1, Jak2, STAT5a/b and Shp2 upon TPO stimulation of *WT* and *Mpig6b*$^{mut}$ MKs. Values are mean ± SD (n = 5). One-way ANOVA with Sidak correction for multiple comparisons. *P < 0.05; **P < 0.01; ***P < 0.001.

**Figure S4. Targeting strategy to generate a novel *Mpig6b*$^{-/-}$ mouse line.** (**a**) A new *Mpig6b*$^{-/-}$ mouse strain lacking exon1 and part of exon2 has been generated by CRISPR/Cas9-based Extreme Genome Editing technology (Biocytogen Inc., China). The two sgRNAs were designed to generate a less than 500 base pair chromosomal deletion in the *Mpig6b* gene. (**b**) Absence of G6b-B was validated in *Mpig6b*$^{-/-}$ platelets by flow cytometry. Platelet count (**c**) and volume (**d**) in 6-week-old female *WT* and *Mpig6b*$^{mut}$ mice were assessed by an automated blood cell analyzer. Values are mean ± SD. Unpaired, two-tailed Student's t-test. *P < 0.05.

**Table S1. Altered GP surface exposure in *Mpig6b*$^{mut}$ platelets.** Fluorescence intensity of major glycoproteins was assessed by flow cytometry. Values are mean ± SD (n = 4). Unpaired, two-tailed Student's t-test. *P < 0.05; **P < 0.01.

**Movie S1. Two-photon intravital microscopy of BM MKs lining vessel sinusoids in 5-week-old *WT* mice.** MKs were visualized with an anti-GPIX antibody derivative coupled to AlexaF546. Vessel lumen is shown in red using BSA-FITC as well as an anti-CD105 antibody coupled to AlexaF488. 20x objective; Frame: 5514.83 ms; 1029 x1029 pixels. Scale bar: 50 µm.

**Movie S2. MK clustering at sinusoidal vessels in 5-week-old male *Mpig6b*$^{mut}$ mice.** MKs were stained in green using anti-GPIX-AlexaF546. Vessels are shown in red (BSA-FITC and anti-CD105-AlexaF488). 20x objective; Frame: 5450.52 ms; 1017x1017 pixels. Scale bar: 50 µm.

**Movie S3. Fragmented MKs and markedly reduced platelet numbers in 5-week-old female *Mpig6b*$^{mut}$ mice.** Vessel staining was performed using BSA-FITC as well as an anti-CD105 antibody coupled to AlexaF488. MKs are stained for GPIX (AlexaF546). 20x objective; Frame: 2900.9 ms; 1017x1017 pixels. Scale bar: 50 µm.

| | *WT* | | *Mpig6b*$^{mut}$ ♂ | | *Mpig6b*$^{mut}$ ♀ | |
|---|---|---|---|---|---|---|
| | mean | SD | mean | SD | mean | SD |
| GPIbα | 491,25 | ±68,3 | 410,25 | ±6,4 | 435,75 | ±27,37 |
| GPIX | 603,75 | ±57,14 | 638,25 | ±41,63 | 635,5 | ±61,26 |
| GPV | 351,25 | ±39,32 | 281 | ±4,32* | 245,5 | ±14,2** |
| CD9 | 1213,25 | ±112,62 | 1004,25 | ±25,38* | 987,5 | ±80,98* |
| GPVI | 69 | ±8,49 | 36,25 | ±1,26** | 36,75 | ±3,2** |
| αIIbβ3 | 634,75 | ±43,02 | 760,75 | ±60,9 | 625,5 | ±66,06 |
| α2 | 71,25 | ±3,77 | 73,75 | ±1,26 | 71,5 | ±1,29 |
| β3 | 328 | ±50,02 | 432,25 | ±22,88 | 375 | ±15,45 |
| CLEC-2 | 187 | ±37,6 | 160 | ±18,81 | 176,25 | ±21,78 |

**Supplemental Table 1**

**a**

| 3 candidates: | prediction: |
|---|---|
| ***Mpig6b*** **(c.404-1G>A)** | **splice acceptor variant (disease-causing)** |
| *Git1* (c.1932C>T) | synonymous variant (p.Leu644Leu) |
| *Prdm1* (c.43-192C>A) | non-pathological intronic variant |

**b**

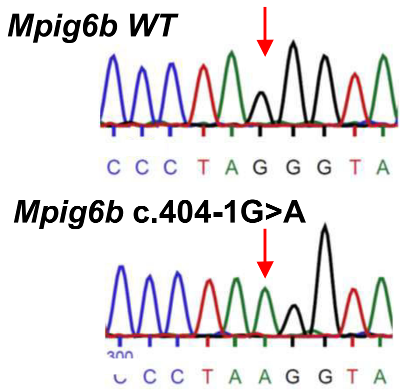


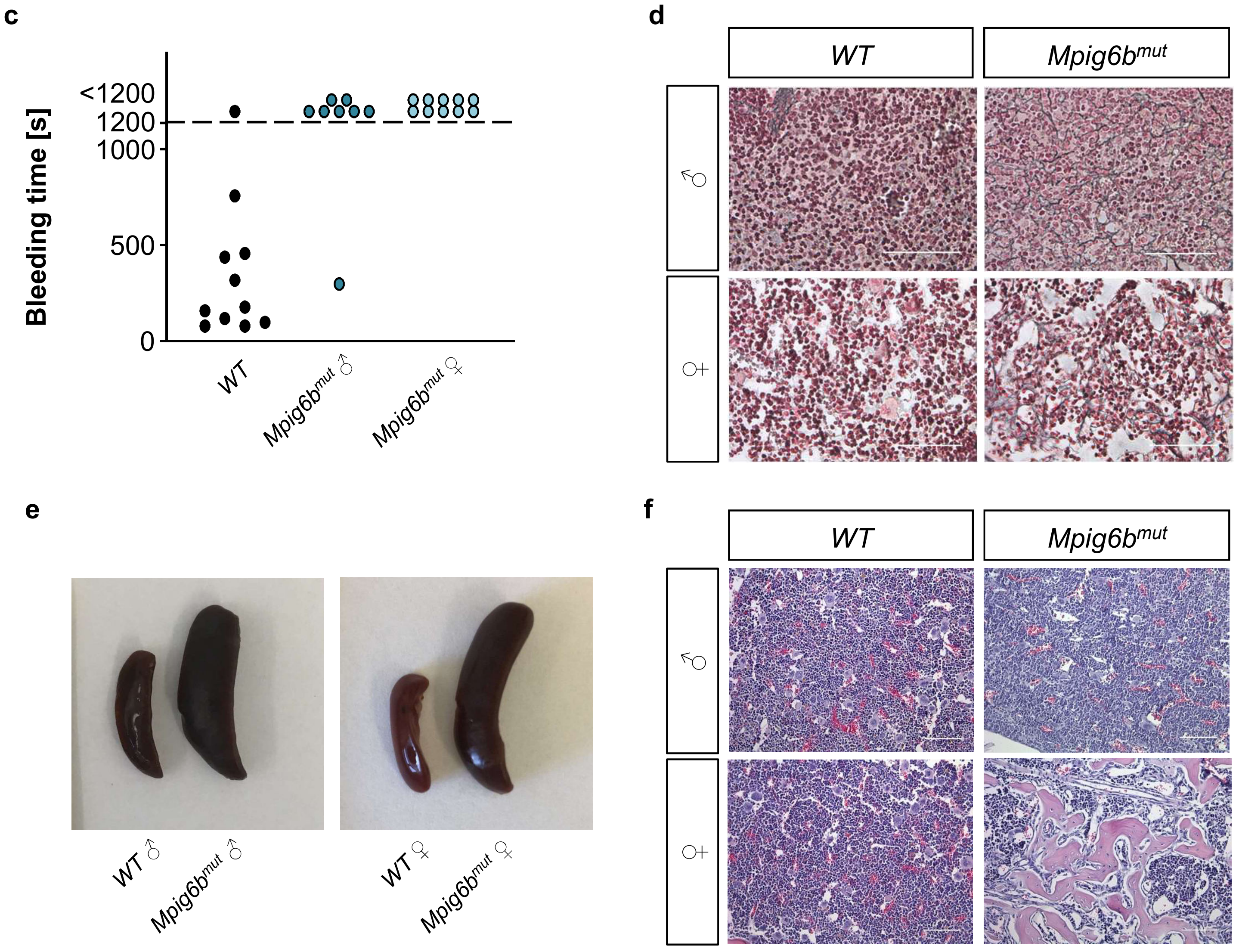


**Supplemental Figure 1**

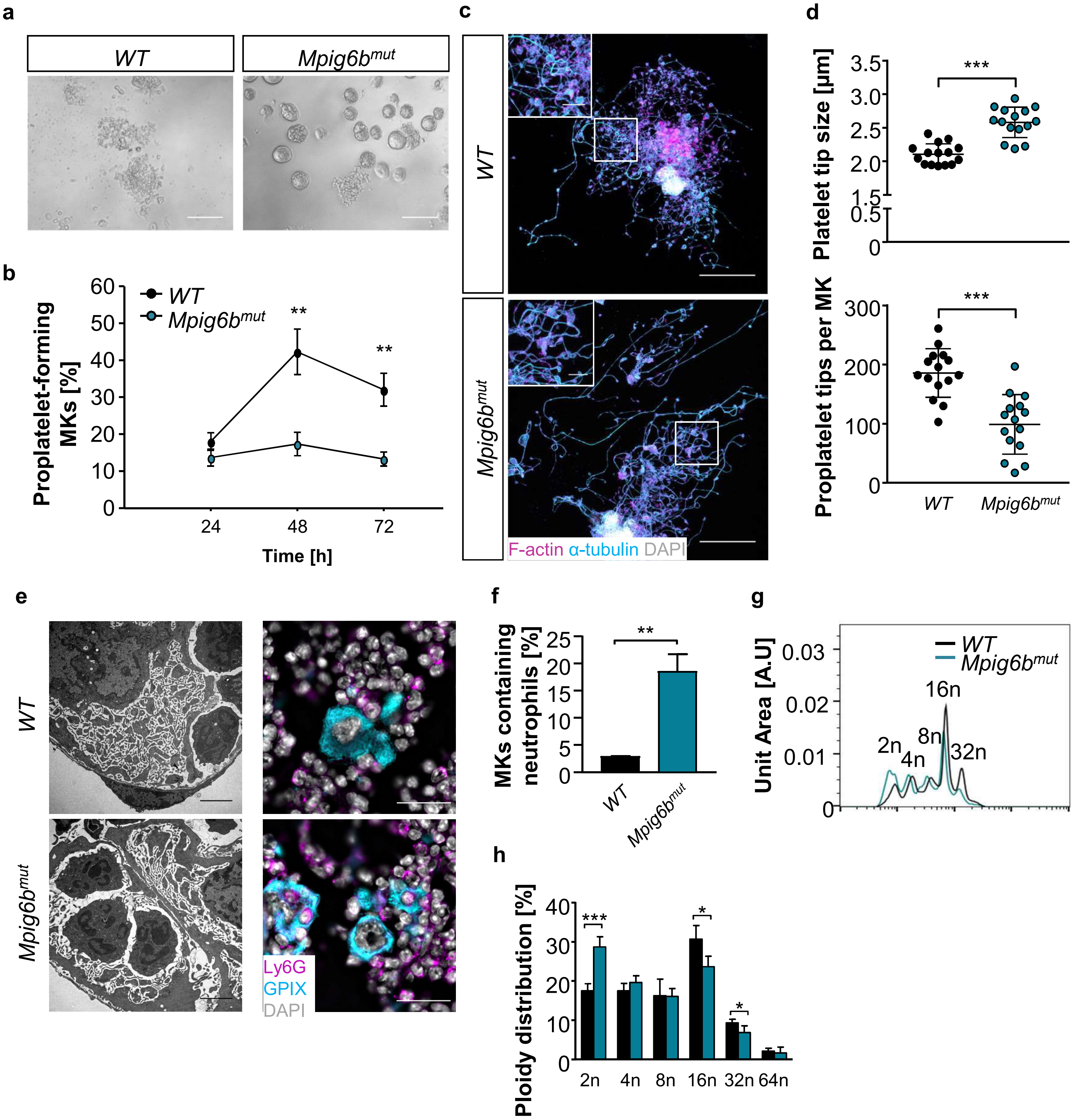


Supplemental Figure 2

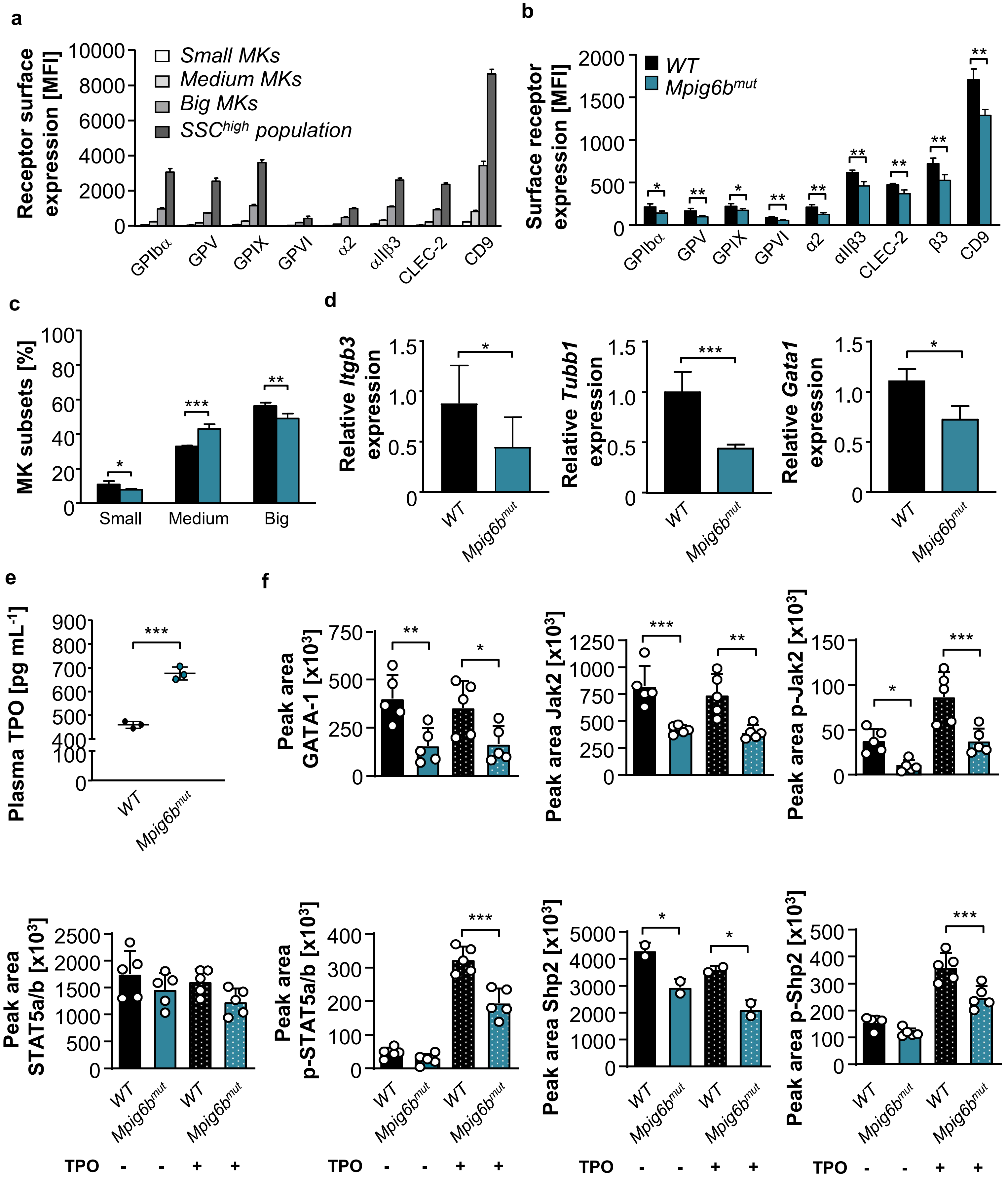


**Supplemental Figure 3**

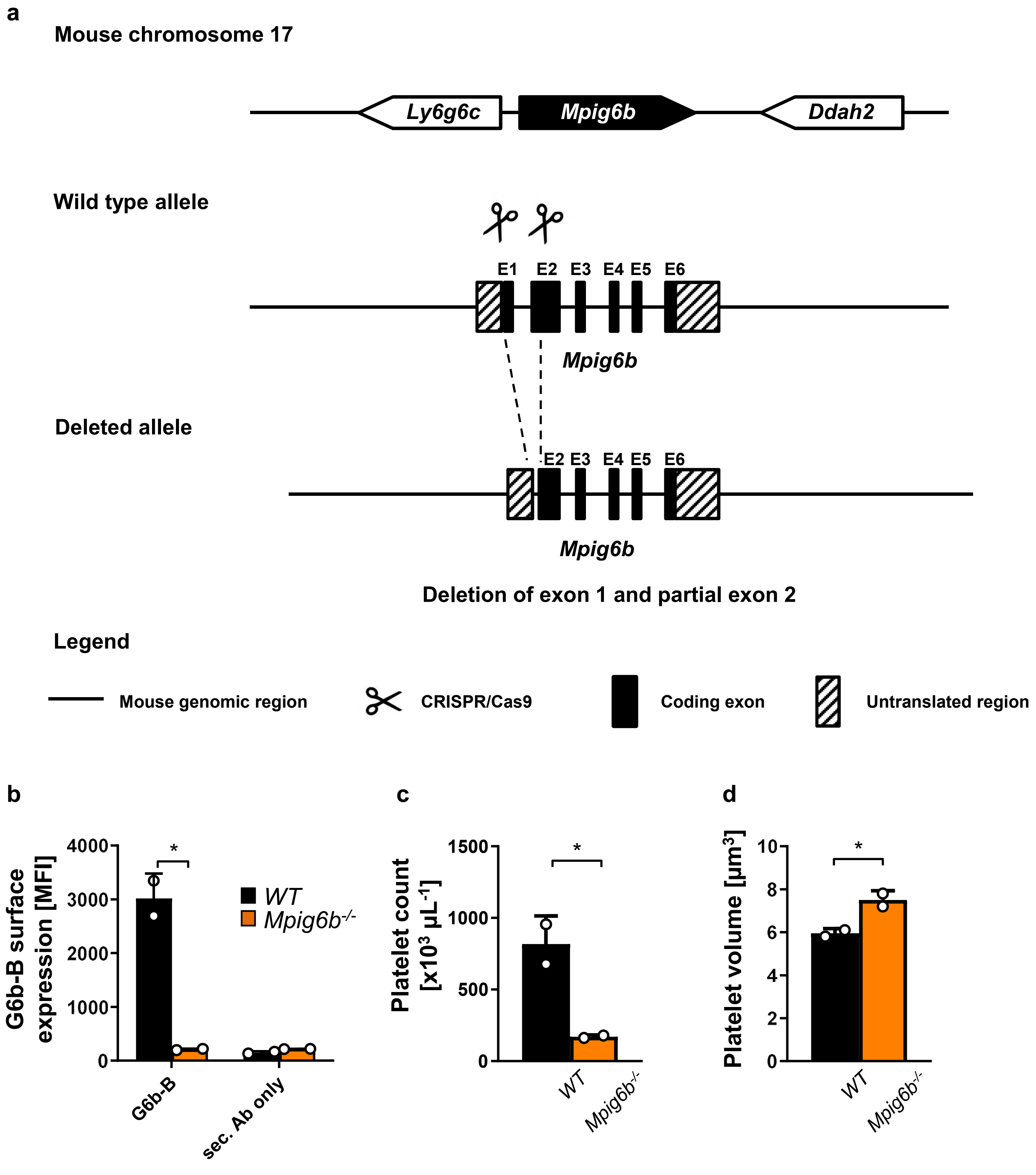


Supplemental Figure 4
